# Orofacial movements: Individuality and stereotypy when mice move a single whisker to touch

**DOI:** 10.1101/2022.10.03.510596

**Authors:** Marcel Staab, Keisuke Sehara, Nora-Laurine Bahr, Sina E. Dominiak, Matthew E. Larkum, Robert N. S. Sachdev

**Author notes:** Corresponding authors: Robert Sachdev (RS); Keisuke Sehara (KS). These authors contributed equally to this work (Marcel Staab, Keisuke Sehara). KS: Department of Physiology, Graduate School of Medicine, The University of Tokyo, Tokyo 113-0033, Japan. SD: Sussex Neuroscience, School of Life Sciences, University of Sussex, Brighton BN1 9QG, UK.

## Abstract

A key function of the brain is to move the body through a rich, complex environment. When rodents engage with their environment, they move their whiskers as they extract tactile information. Even though the study of whisking has a long history, the details of individual whisker movements bilaterally, of nose movement, of stereotypy and variability in an active whisking to touch task are unknown. Here we trained head fixed mice in a simple go-cue task to move a whisker on one side of the face to touch a sensor and tracked facial movements. Our analysis shows that mice specifically control movement of the whisker they use to touch and that as they move their whiskers, they move their nose and apply forces on the head-post in a manner that reflects the behavioral epoch, i.e. whether go cue triggered movement had begun, or a whisker was touching the sensor. Importantly, mice control the setpoint, amplitude and frequency of movement of whiskers bilaterally and individually. Additionally, even though mice achieved the goal of the task -- to touch the sensor within 2 seconds -- how they coordinated movement of the nose and forces on head post with movement of individual whiskers was stereotyped and related to the distance they needed to move a whisker to touch the sensor. Our work shows how stereotyped mouse behavior can be, and it emphasizes both the level of fine motor control mice can exert over individual whiskers and the extent of facial movements in a goal-directed whisking-to-touch task.

**Significance:** Recent work shows that facial movements are reflected in the activity of a surprisingly large number of brain areas. But what aspects of the face do mice move when they move a whisker to actively touch an object? Our work shows that while mice control the movement of a whisker they use to touch, they also move their nose, apply forces on the head-post and move adjacent whiskers and whiskers on the other side of the face. Additionally, our analysis shows that from day-to-day, this behavior can be surprisingly stereotyped, and that small changes in how far mice move a whisker during tactile behavior fundamentally changes the relationship between the movement of a single whisker and other facial movements.

## Introduction

Over the course of evolution, and over the course of the lifetime of an animal, the motor sequences used to achieve a particular goal are refined and movement of many parts of the body are linked together into a whole that is optimal for achieving a particular goal. While we know that most natural behavior is multi-modal (Krakauer et al., 2017), involving many sensory and motor modalities, how various parts of the body move to generate a particular behavior, how variability in the movement of any part of the body affects movement of other parts of the body, and generates activity in a variety of sensory systems is only now beginning to be studied (Churchland, Afshar & Shenoy, 2006; Ghazanfar & Schroeder, 2006; Kelso, 2009; Krakauer et al., 2017; McElvain et al., 2018; Murakami & Mainen, 2015; Waschke et al., 2021).

Even in a model system like the rodent whisker system, where the goal of whisking is first and foremost to simply palpate surfaces and use tactile input to guide the animal around its environment, we are only on the cusp of understanding how rodents coordinate their facial movements (Vincent, 1912; Carvell & Simons, 1990; Hartmann, 2011, 2001; Sachdev, Sato & Ebner, 2002; Berg & Kleinfeld, 2003; Knutsen, Pietr & Ahissar, 2006; Grant et al., 2009; Haidarliu et al., 2010, 2012, 2015; Deschênes, Moore & Kleinfeld, 2012; McElvain et al., 2018; Adibi, 2019). For example, recent work has revealed that facial expressions in mice can convey emotions like pain, disgust or joy (Langford et al., 2010; Finlayson et al., 2016; Dolensek et al., 2020), and that facial movements can explain variance in single unit activity in surprisingly widespread areas of the brain (Avitan & Stringer, 2022; Stringer et al., 2019; Zagha et al., 2022; Steinmetz et al., 2019; Musall et al., 2019).

Traditionally, work in the whisker system focused on simple organizational features. Firstly, input from each whisker forms a labelled line from the trigeminal brainstem to cortex (van der Loos & Woolsey, 1973). Secondly, while whiskers are associated with the whisker pad that can be moved by facial muscles, each whisker has a single muscle slung around it (Dörfl, 1982; Wineski, 1983, 1985; Haidarliu et al., 2010); thirdly, whisking is associated with active behavior that can be directional, reflecting direction of movement of the animal (Knutsen, Derdikman & Ahissar, 2005; Towal & Hartmann, 2006; Grant, Mitchinson & Prescott, 2012; Grant, Sperber & Prescott, 2012; Arkley et al., 2014; Saraf-Sinik, Assa & Ahissar, 2015; Schroeder & Ritt, 2016; Dominiak et al., 2019; Bergmann et al., 2022); fourth, whisking and breathing are linked together, by a central pattern generator in the brain stem (Welker, 1964; Cao et al., 2012; Moore et al., 2013; Moore, Kleinfeld & Wang, 2014; McElvain et al., 2018; Tantirigama et al., 2020); and finally, while whisking is central for navigation, mice also move their eyes as they navigate and they coordinate their movement with the movement of their whiskers, nose and eyes (Dominiak et al., 2019; Bergmann et al., 2022). Despite the relative sensory “isolation” of each whisker (but see, Severson et al. 2017; Severson et al. 2019) and the potential for independent control of movement of each whisker (Sachdev, Sato & Ebner, 2002; Grant et al., 2009; Haidarliu et al., 2010), the ability of rodents to control the movement of individual whiskers and to coordinate this motion, with movement of other aspects of the face remains largely unexplored.

Here we begin the task of specifically measuring the coordination, variability and individuality that defines mouse whisking behavior, in a simple go-cue triggered whisking to touch task. Our work shows that in this task mice move individual whiskers distinctly on the two sides of the face and they coordinate motion of the nose and forces on the head post in a specific manner. This work also shows that from day to day the individual movement patterns of each mouse can be measurably stereotyped in a goal directed manner.

## Results

### Whisking, nose motion and forces on head post

Sixteen mice were trained to use the right side C2 whisker to contact a sensor within 2 second of an auditory cue (Fig 1). In the course of 5964 correct trials over 84 sessions, the movement of the nose, and the painted C1 and C2 whiskers bilaterally were tracked as mice whisked to touch the sensor, waited for the reward, and licked (Fig 1A). The maximum intensity projection for a single session (n=40 trials) from one mouse shows the extent of movement of two whiskers on each side of the face, and the nose on all successful trials (Fig 1B). Additionally, a strain gauge was used to measure the up and down forces mice applied on the head post (Fig 1C). The movement / position of whiskers and nose, and the forces mice applied on the head post, on each trial, were converted into raster displays reflecting the position and movement of whiskers and side to side motion of the nose, and the extent of the forces applied on the head post in each trial (Fig 1D). This display shows the timing and extent of whisker and nose motion, and the dynamics of the coordination between movements and the forces on the head post on every trial. The median position for the 40 trials in a session are plotted below each raster plot.

**Figure 1.**
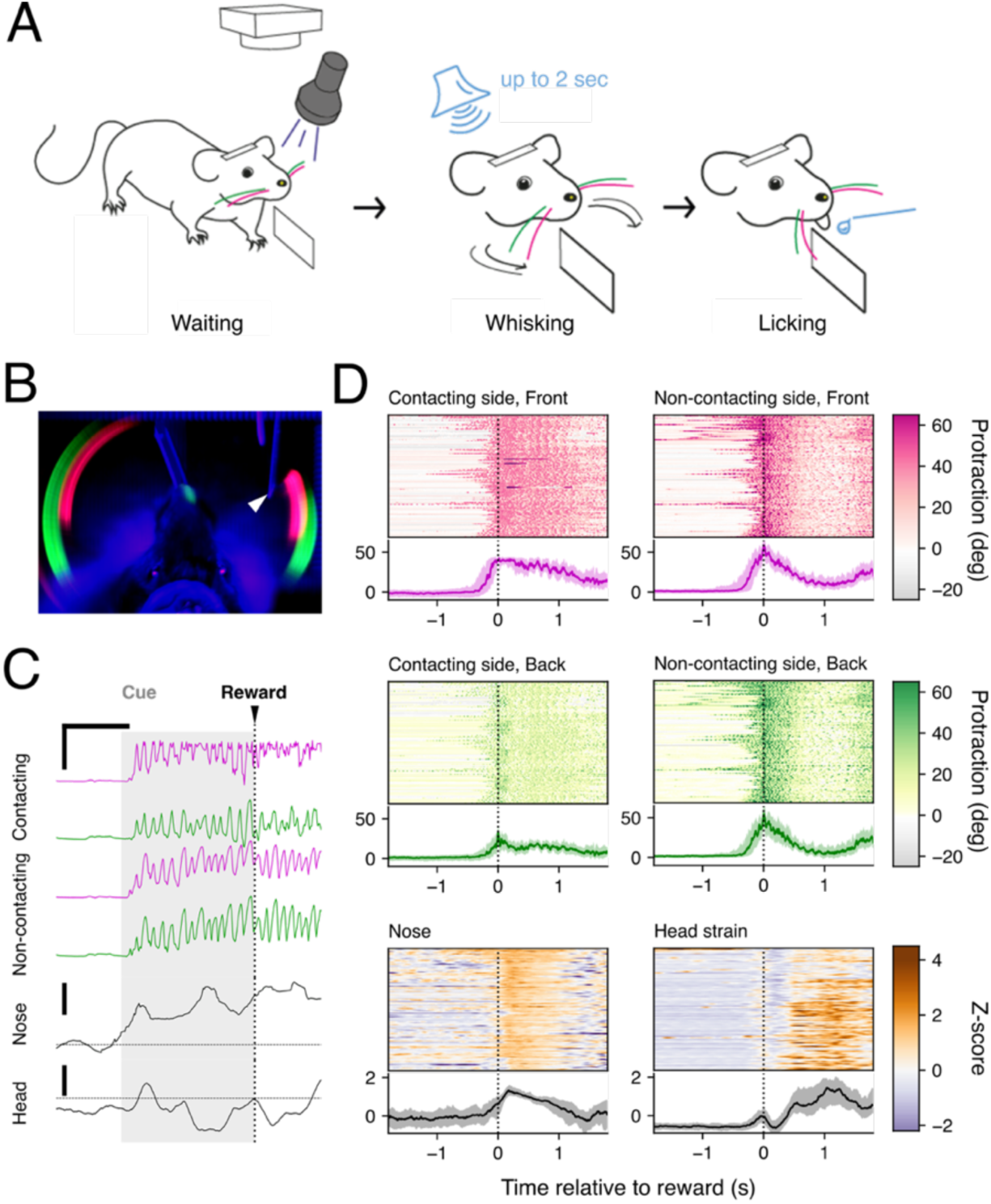
Bilateral whisker and nose motion, and strain on a head post during whisking to touch. A. Schematic of the task. Mice were head-fixed, and their C2 (magenta) and C1 (green) whiskers were dabbed with UV paint. In response to the auditory cue (up to 2 seconds), mice moved the C2 whisker on the right side of the face to contact the iezo sensor in front of their face (middle) to obtain water reward (right). B. Top-view maximal intensity projection of a representative session. Whisker trajectories for the entire session are visible as magenta and green arcs around the animal. The arrowhead on the right side indicates the position of the contact sensor. C. Tracking whiskers and nose position and head post forces in a single trial. Top four traces show the whisker positions in the course of a trial on the **contacting** side and the **non-contacting** side. The traces in magenta represent positions of the two C2 whiskers (the front whisker) whereas those in green represent positions of the C1 whisker (the whisker in back). Mediolateral side to side positions of the nose in the course of the same set of trials, and the values of the up and down head strain sensor are shown at bottom. The horizontal scale bar indicates 500 ms. The vertical scale bars, 60° for whiskers and 1 Z-value for nose position and head strain. D. Raster plots of tracked behavior of a single representative session. The protraction angles of individual whiskers, and the Z-score values of the nose position and the head strain signal, from individual trials are color-coded at the top of each panel. Thick lines at the bottom represent session medians, and the shaded areas represent 25–75 percentile ranges during the session.

This representative session reveals that, even though mice are highly trained to move the C2 whisker on the right side into contact with the sensor, and they reliably contact the sensor with just this one whisker, the mouse invariably moves all the tracked whiskers bilaterally on every trial. Even though mice move their whiskers bilaterally on every trial, the movement of the whiskers on the two sides of the face -- the contacting side and the non-contacting side -- were not identical. On the non-contacting side, the C1 and C2 whiskers move together, and show similar movement patterns: fast protractions and large movements. On the contacting side, the C1 whisker moves much less than the C2, and both whiskers show a different pattern of movement than on the non-contacting side. Part of the differences in motion on the two sides of the face simply reflect the presence of the contact sensor that blocks large movements of the front, C2 whisker, but part of the movement especially of the C1 whisker in the back reflects the independent control of this whisker. These analyses also show the trial by trial and session-long stereotypy of the side-to-side nose motion and the head strain, as the mouse touch the contact sensor and lick reward.

### Side-to-side differences in multi-whisker behavior

To elucidate the behavioral strategies mice used in this task, we first examined the movement of individual whiskers during the task in more detail (Figs 2 and 3). The positions of whiskers were converted into an animal-centric coordinate system, using the position of the eyes (Fig 2A). Whisker movement was converted into angular motion and separated into whisking setpoint and amplitude by examining the envelope of the whisking motion (Fig 2B).

**Figure 2.**
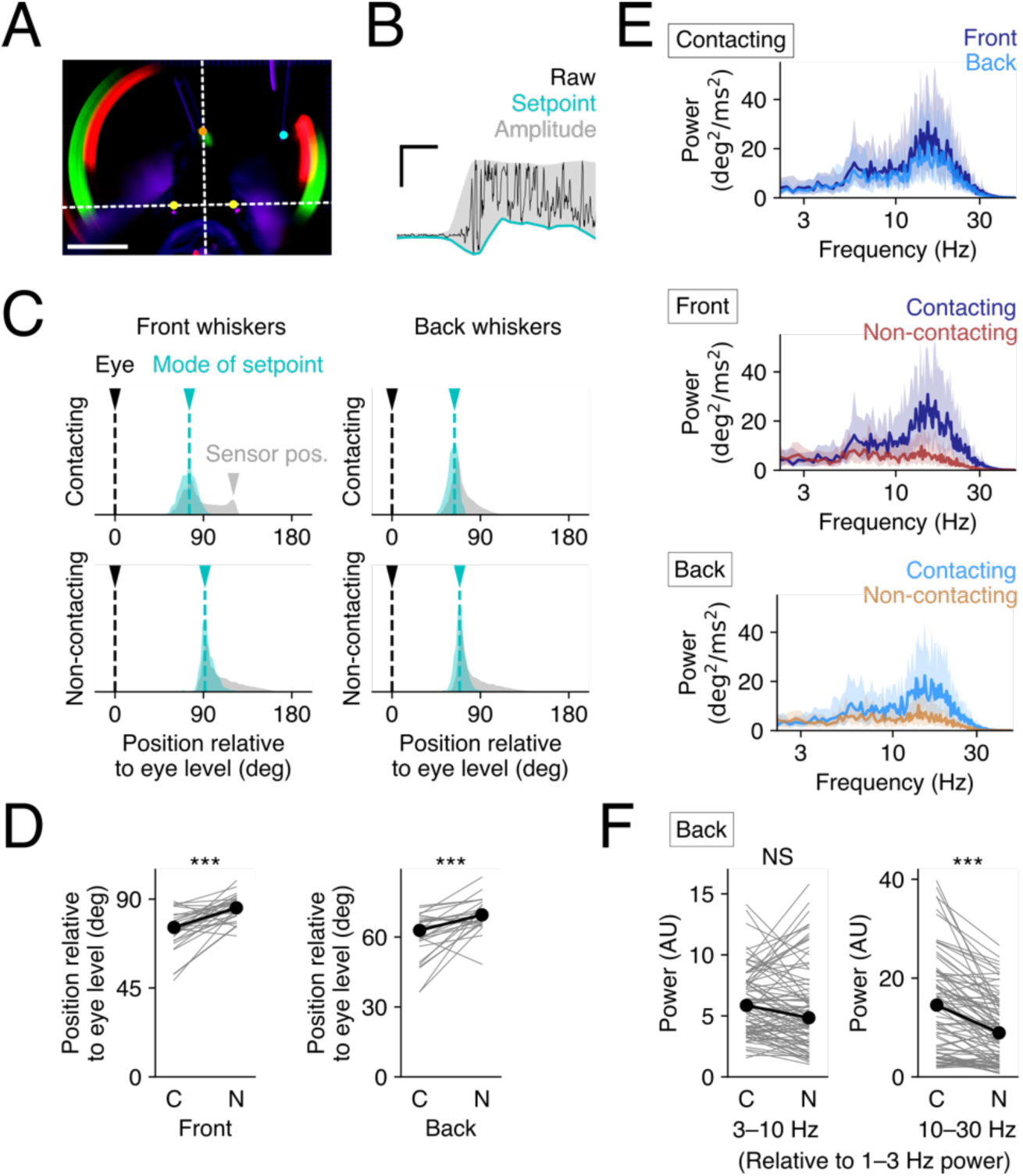
Control of single whisker movement. A. Comparison of resting whisker positions relative to the level of the eyes. The position of the eyes was manually annotated (yellow) -- and the white horizontal dotted line denoting the position of the eyes -- was used as coordinate system for setting the angles for the whiskers. Eye position was set as zero degrees. Scale bar, 10 mm. B. Schematic for estimation of setpoints and amplitude from raw whisker traces (black). A single representative trial is shown as an example where the lower envelope (cyan) was defined as the setpoint, and the difference between the lower and the upper envelope (gray) was defined as the amplitude of whisking. C. Distribution of setpoints positions for whiskers on the contacting and the non-contacting sides. The mode in the histogram of setpoints (cyan) during each session was defined as the resting position of the whisker during the session, the gray shaded areas shows the distribution of whisker positions in the whole the session. Black arrowheads mark the eye position, cyan arrowheads mark the resting whisker position, and gray arrowhead (top left panel) marks the location of the contact sensor (gray). D. Statistical comparison of resting positions on the two sides of the face. The C2 and C1 whiskers on the contact side have significantly different resting positions than the C2 and C1 whiskers on the non-contact side. C, contacting side; N, non-contacting side; ***p<0.001, Wilcoxon signed rank test. N=29 sessions from 6 animals. E. Power spectra of whisker speeds. The results of a single representative session show the distribution of whisking frequencies in the course of whisking to touch. Note that whiskers on the contacting side, both the front and the back show much higher frequency components --at ∼25 Hz, then the whiskers on the non-contacting side. Thick lines represent median of the trials of the session, and the shaded areas correspond to 25–75 percentiles. F. Comparison of the whisking power of low (3–10 Hz, left) and high (10–30 Hz, right) frequency bands of whisking speed. The values are normalized relative to the power of the 1–3 Hz band. ***p<0.001; NS, p>0.05, Wilcoxon signed rank test. N=85 sessions from 16 animals.

**Video 1.**
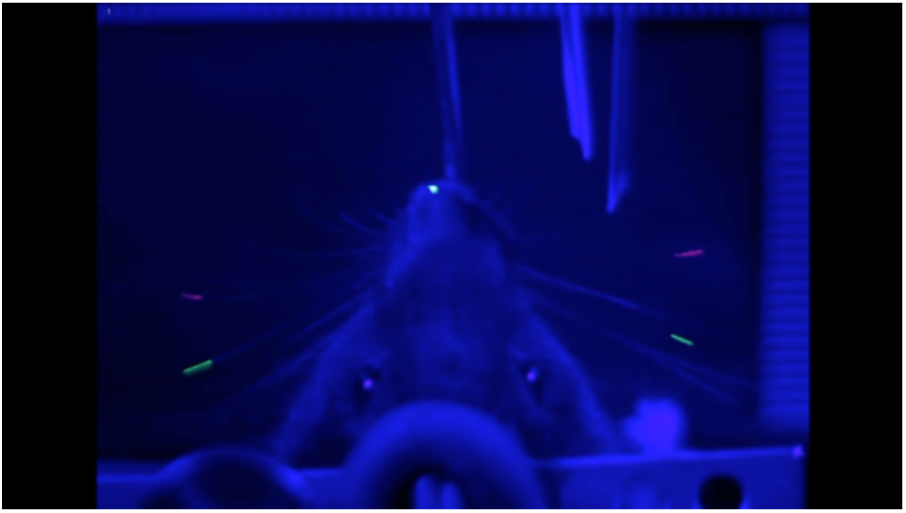
Contact sensor on right, positioned near C2 whisker.

**Video 2.**
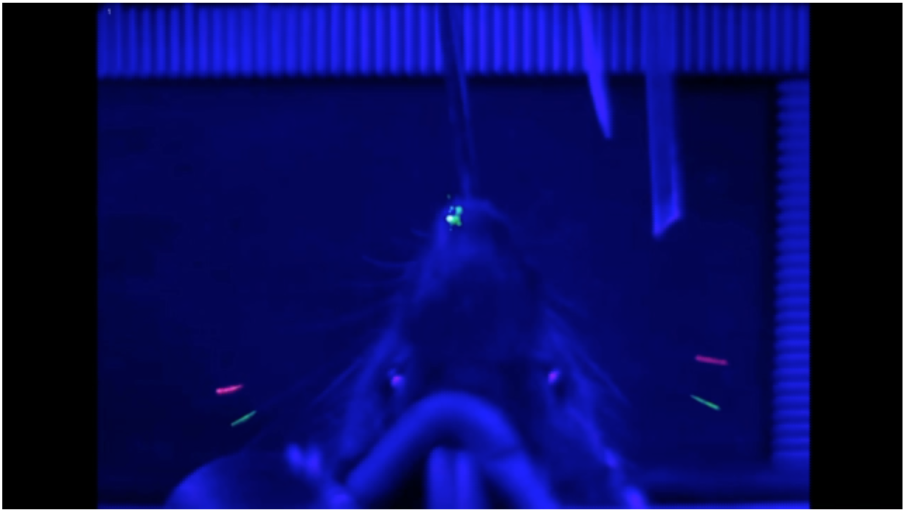
Contact sensor on right, C2 whisker retracted.

The maximal intensity projections of tracked whiskers (Fig 2A) indicated that the resting position of the tracked whiskers on the two sides of the face were different. Examination of individuals of mice engaged in the task supported this possibility (1-2). To quantify this difference, the mode of the setpoint for each whisker during a given behavioral session was calculated and considered to be the resting position (Fig 2C). This analysis shows that the resting position of the whiskers on the non-contacting-side (relative to the level of the eyes) was more protracted than the resting position of the whiskers on the contacting side (Fig 2C). Comparison across multiple sessions confirmed that the resting whisker position on the non-contacting side was significantly more protracted than on the contacting side (Fig 2D; front whiskers, p=0.0001***; back whiskers, p=0.0001***; Wilcoxon signed rank test, N=29 sessions of 6 animals).

Next, we examined the frequency components of whisking on the two sides of the face (Fig 2E-F). The power spectra of whisking traces for a single session on the contacting side for the front and back whiskers showed a bimodal / skewed distribution, with a peak at ∼25 Hz. Note that the peak at 25 Hz does not result from contact. The C1 whisker (back) adjacent to the contacting whisker does not contact the sensor at all (Fig 2E, light blue traces). The histogram of positions for the C1/back whisker was not affected by the contact sensor (Fig 2C, top right, gray area). This high frequency whisking band was prominent on the contacting side (Fig 2E).

To compare the frequency components detected across all sessions from all animals, we normalized the power of the low (3–10 Hz) and the high (10–30 Hz) frequency bands relative to the baseline (1–3 Hz) power. This analysis indicates that there was significantly more power at higher frequencies of whisking on the contacting side than on the non-contacting side (Fig 2F, right) (3–10 Hz, p=0.0623, NS; 10–30 Hz, p=0.0001***; Wilcoxon signed rank test, N=85 sessions of 16 animals). There were no significant differences in side-to-side power at lower frequencies of whisking (Fig. 2F, left).

Together, these results highlight the side-to-side asymmetry in whisking. Compared to the whiskers on the non-contacting side, the contacting side whiskers tended to be retracted during the task and tended to move with more vigor at higher frequencies.

### Behavioral phase-dependent single-whisker movement dynamics

Next, we examined how changes in setpoints and whisking amplitudes characterize the whisking behavior of the mice during the task (Fig 3). Single session data illustrate the distinct setpoint of individual whiskers (Fig 3A). Note that the setpoint data shown here was defined as a deviation of the whisker position from the resting whisker position during the session. This definition avoids the bias introduced by day-to-day or whisker-to-whisker (C2, C1 positions) differences in resting positions. We see that on average, the setpoint of the whisker used to touch the sensor was more protracted than the setpoint of the adjacent whisker (Fig 3A, left). By comparison, whiskers on the non-contacting side moved almost identically, with their setpoints being similar to each other (Fig 3A, right).

**Figure 3.**
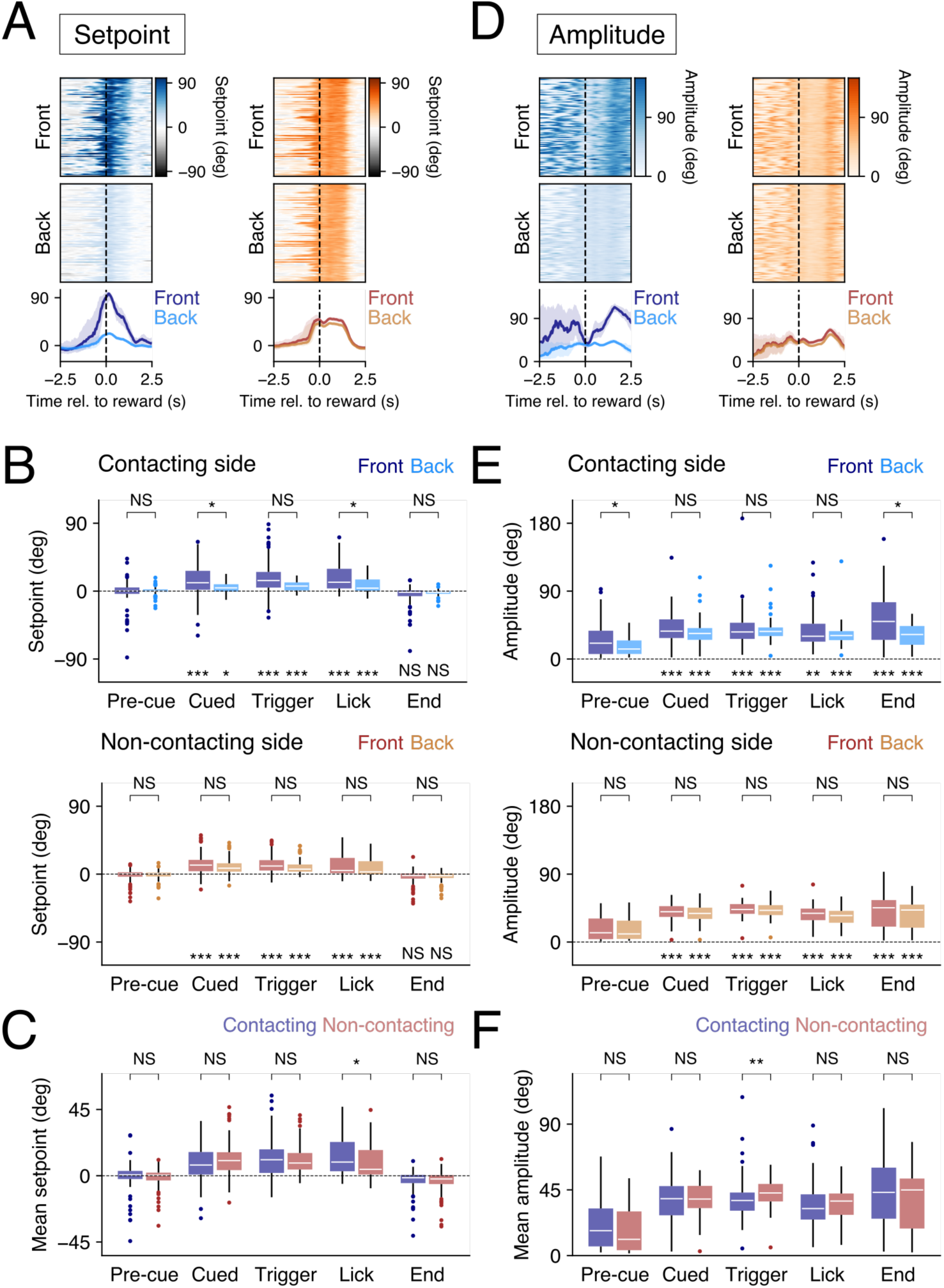
Behavioral phase-dependent multi-whisker dynamics. A. Whisker-to-whisker variability of setpoint changes in a single representative session. Raster plots represent estimated setpoints of front (top panels) and back (middle panels) whiskers on the contacting side (left panels) and on the non-contacting side (right panels) for individual trials. The bottom panels show session median traces for the whiskers. Setpoint protraction tended to be larger for the front whiskers than the back whiskers, and the tendency was more visible on the contacting side. B. Comparison of setpoints of adjacent whiskers on contacting and non-contacting side. All the whiskers protracted significantly during the cued, triggering, and licking period compared to during the pre-cue period, but the front whisker only on the contacting side was significantly more protracted than the adjacent whisker; the deviations in setpoint between the front and the back whiskers increased during the cued and triggering periods (top panel). There were no significant differences between adjacent whiskers on the non-contacting side. Session-median setpoints during each of the five behavioral time windows are shown as a Tukey’s box plot. Results from both contacting (blue) and non-contacting sides (red) are shown. **N=82 sessions for the cued time window, and N=85 sessions for the other time windows.** C. Comparison between setpoints of whiskers of contacting and non-contacting side. (D) Whisker-to-whisker changes in amplitude of whisking in a single representative session, in the same manner as in (A). E. Statistics for amplitude shown in the same way as in (C). Mice made larger whisking movements after the cue onset on both sides of the face. The difference in amplitude of whisking of the contacting front whisker and adjacent back whisker was only significant during some of the behavioral epochs. There were no significant differences in amplitude of whisking of adjacent whisker on the non-contacting side (bottom panel). *p<0.05, ***p<0.001, NS, p>0.05, Kruskal–Wallis test followed by *post hoc* Dunn’s pairwise tests with Bonferroni correction**. N=82 sessions for the cued time window, and N=85 sessions for the other time windows. F. Comparison of amplitude of whisking on the contacting and the** non-contacting sides. Same analysis as for setpoint in (C).

To examine the behavioral state dependent changes in whisking in more detail, each trial was divided into distinct 750 ms epochs (Table 1). At cue onset, mice adjusted the setpoint of whiskers bilaterally; they protracted their whiskers and kept them protracted until the licking phase of the trial (Fig 3B, Tables 2 and 3, p<0.05, Kruskal–Wallis test, followed by Dunn’s *post hoc* pairwise tests with Bonferroni correction).

On the contacting side, during the cued and the licking phase of the trial, adjacent whiskers were positioned distinctly, with the front whisker (C2) being kept significantly more protracted than the back whisker (C1) (Fig 3B, top; Table 2, Front vs Back; p<0.05, Dunn’s *post hoc* pairwise tests with Bonferroni correction). On the non-contacting side mice protract their whiskers but there were no significant differences in the positioning of the front and the back whiskers, for any of the behavioral phases examined (Fig 3B, bottom; Table 3, Front vs. Back; p>0.05, Dunn’s *post hoc* pairwise tests with Bonferroni correction).

**Table 1.** Definitions of the behavioral epochs. Each trial was divided into 750 ms epochs defined on the basis of the timing of the auditory cue onset or on the basis of the contact sensor trigger. Note that there were some trials without pre-cue or cued periods. Because the recorded videos contained only 2.5 seconds before the contact trigger, trials were considered not to have the pre-cue period if the auditory cue duration was more than 1.75 seconds. Similarly, the trial was considered not to have the cued period if the cue duration was less than 750 ms.

| Behavioral phase | Start of the phase | End of the phase |
| --- | --- | --- |
| Pre-cue | 750 ms before the cue onset | Cue onset |
| Cued | Cue onset | 750 ms after the cue onset |
| Trigger | 500 ms before the contact trigger | 250 ms after the contact trigger |
| Licking | 250 ms after the contact trigger | 1000 ms after the contact trigger |
| End of trial | 1750 ms after the contact trigger | 2500 ms after the contact trigger |

Note that during the cued epoch, the difference in the whisking setpoint of the adjacent (front-back) whiskers was not likely to arise from side-to-side differences in whisking. In this epoch on average, there were no significant differences between the whisking setpoints on the contacting and non-contacting side whiskers (Fig 3C; pre-cue, p=0.5148, NS; cued, p=0.2371, NS; trigger, p=0.9156, NS; lick, p=0.0318*; end-of-trial, p=0.8664, NS; Mann–Whitney U-test, N=85 sessions from 16 animals). Taken together, these results indicate that, when mice are engaged in the task, they protract their whiskers differentially on the two sides of the face and specifically control the spread between whiskers.

**Table 2.** Pairwise statistics on contacting side whisking setpoints. Pairwise comparisons of whisker setpoints in the different behavioral epochs showing the change in setpoint in the active epochs when compared to the pre-cue epoch. Both the contacting whisker “Front” and the “Back” adjacent whisker were positioned at distinct setpoints during behaviors: cue, trigger and licking. There were significant differences in setpoints, for contacting (front) and adjacent whisker (back) during the cue and lick epochs. Analysis was performed using a Kruskal–Wallis test, followed by Dunn’s *post hoc* pairwise tests with Bonferroni correction. For each session, the median values of whisking setpoint, in each behavioral phase, for each whisker were computed and the position of the whisker in each behavioral epoch were compared to position in the pre-cue epoch. The omnibus p-value was p=0.0000. Each row on the table corresponds to a *post hoc* pairwise testing**. N=85 sessions of 16 animals**.

| Category | Comparison | P-value |
| --- | --- | --- |
| Contacting side setpoint, Front | Pre-cue vs. Cued | 0.0000*** |
|  | Pre-cue vs. Trigger | 0.0000*** |
|  | Pre-cue vs. Lick | 0.0000*** |
|  | Pre-cue vs. End-of-trial | 0.1463, NS |
| Contacting side setpoint, Back | Pre-cue vs. Cued | 0.0456* |
|  | Pre-cue vs. Trigger | 0.0001*** |
|  | Pre-cue vs. Lick | 0.0002*** |
|  | Pre-cue vs. End-of-trial | 0.1871, NS |
| Contacting side setpoint, Front vs. Back | Pre-cue | 1.0000, NS |
|  | Cued | 0.0219* |
|  | Trigger | 0.0627, NS |
|  | Lick | 0.0298* |
|  | End-of-trial | 1.0000, NS |

**Table 3.** Pairwise statistics on non-contacting side whisking setpoints. Pairwise comparisons of whisker setpoints showing the change in starting position / set point in the active epochs when compared to the pre-cue epoch. Mice actively moved, positioned whiskers on the non-contact side in the course of the task, but adjacent whiskers on this side of the face moved and were positioned similarly, there were no significant differences in the front and back whiskers position in the different behavioral epochs. Analysis was performed using a Kruskal–Wallis test, followed by Dunn’s *post hoc* pairwise tests with Bonferroni correction. For each session, the median values of whisking setpoint, in each behavioral phase, for each whisker were computed and the position of the whisker in each behavioral epoch were compared to position in the pre-cue epoch. The omnibus p-value was p=0.0000. Each row on the table corresponds to a *post hoc* pairwise testing. **N=85 sessions of 16 animals.**

| Category | Comparison | P-value |
| --- | --- | --- |
| Non-contacting side setpoint, Front | Pre-cue vs. Cued | 0.0000*** |
|  | Pre-cue vs. Trigger | 0.0000*** |
|  | Pre-cue vs. Lick | 0.0000*** |
|  | Pre-cue vs. End-of-trial | 0.4933, NS |
| Non-contacting side setpoint, Back | Pre-cue vs. Cued | 0.0000*** |
|  | Pre-cue vs. Trigger | 0.0000*** |
|  | Pre-cue vs. Lick | 0.0000*** |
|  | Pre-cue vs. End-of-trial | 0.3723, NS |
| Non-contacting side setpoint, Front vs. Back | Pre-cue | 1.0000, NS |
|  | Cued | 1.0000, NS |
|  | Trigger | 1.0000, NS |
|  | Lick | 1.0000, NS |
|  | End-of-trial | 1.0000, NS |

Next, we examined whisking amplitude. Analysis of average whisking amplitude across sessions demonstrated that when compared to the pre-cue period, it increased significantly on both sides of the face from the onset of the auditory cue until the end of trial (Fig 3D & 3E; Tables 4 and 5; p<0.001, Kruskal–Wallis test, followed by Dunn’s post hoc pairwise tests with Bonferroni correction). Additionally, there were behavioral epoch dependent significant differences in amplitude of whisking for adjacent whiskers on the contacting side. During both the pre-cue and the end-of-trial phases, the whisking amplitude on the contacting side for the front, C2 whisker was significantly larger than that of the adjacent, back whisker, (Table 4, Front vs. Back). There were no significant differences in amplitude of whisking of the two whiskers on the non-contacting side (Table 5, Front vs. Back).

Whisking amplitudes on the two sides of the face did not differ significantly between the contacting and non-contacting sides, except in the contact trigger phase (Fig 3F) (pre-cue, p=0.0854, NS; cued, 0.7836, NS; trigger, 0.0023**; lick, 0.3082, NS; end-of-trial, 0.3156, NS; Mann–Whitney U-test, N=85 sessions of 16 animals). The difference during the contact trigger phase is likely to be related to the presence of the contact sensor which blocks motion of the front whisker on the contacting side.

**Table 4.**
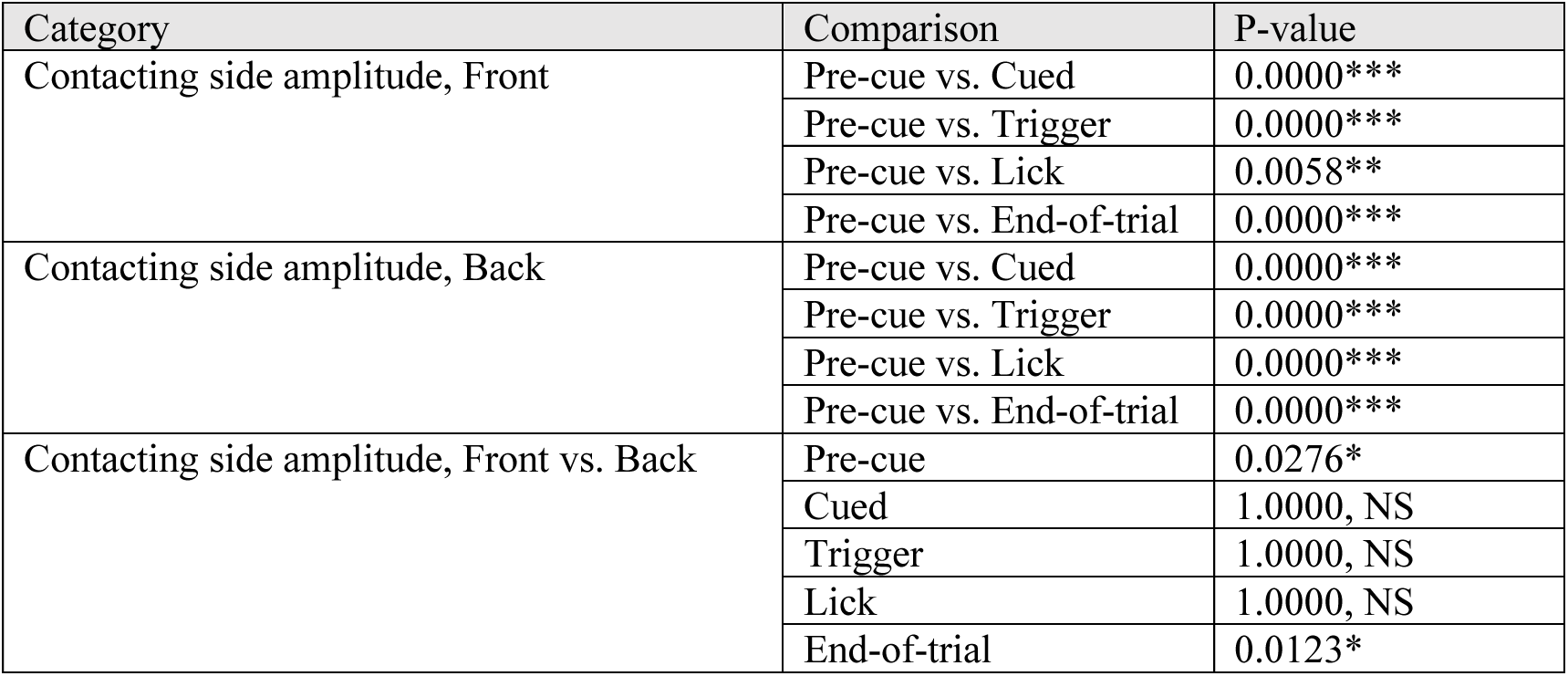
Pairwise statistics on contacting side whisking amplitude. Pairwise comparisons of whisking amplitude in the different behavioral epochs show that amplitude of movement of both whiskers were significantly different from the resting “pre-cue epoch” whisking amplitude. There were some significant differences in whisking amplitude for the two adjacent whiskers, primarily in the Statistical analysis were performed using a Kruskal–Wallis test, followed by Dunn’s *post hoc* pairwise tests with Bonferroni correction. For each session, the median values of whisking amplitude, in each behavioral phase, for each whisker were computed and the amplitude of whisker movement in each behavioral epoch was compared to amplitude in the pre-cue epoch. The omnibus p-value was p=0.0000. Each row on the table corresponds to a *post hoc* pairwise testing**. N=85 sessions of 16 animals.**

Taken together, our analysis indicates that mice exert control over the movement of individual adjacent whiskers on each side of the face and they display distinct movement patterns on the two sides of the face. These differences in individual whisker positions primarily reflect the effect on the setpoint. When mice use a whisker to touch the sensor, the spread of whiskers -- the distance between adjacent whiskers -- changes, and the whiskers spread apart, but only on the side actively used in this tactile behavior.

**Table 5.** Pairwise statistics on non-contacting side whisking amplitude. Pairwise comparisons of whisking amplitude in the different behavioral epochs show that amplitude of movement of both whiskers were significantly different from the resting “pre-cue epoch” whisking amplitude. Statistical analysis a Kruskal–Wallis test, followed by Dunn’s *post hoc* pairwise tests with Bonferroni correction. For each session, the median values of whisking amplitude, in each behavioral phase, for each whisker were computed and the amplitude of whisker movement in each behavioral epoch was compared to amplitude in the pre-cue epoch. The omnibus p-value was p=0.0000. Each row on the table corresponds to a *post hoc* pairwise testing. **N=85 sessions of 16 animals.**

| Category | Comparison | P-value |
| --- | --- | --- |
| Non-contacting side amplitude, Front | Pre-cue vs. Cued | 0.0000*** |
|  | Pre-cue vs. Trigger | 0.0000*** |
|  | Pre-cue vs. Lick | 0.0000*** |
|  | Pre-cue vs. End-of-trial | 0.0000*** |
| Non-contacting side amplitude, Back | Pre-cue vs. Cued | 0.0000*** |
|  | Pre-cue vs. Trigger | 0.0000*** |
|  | Pre-cue vs. Lick | 0.0000*** |
|  | Pre-cue vs. End-of-trial | 0.0000*** |
| Non-contacting side amplitude, Front vs. Back | Pre-cue | 1.0000, NS |
|  | Cued | 1.0000, NS |
|  | Trigger | 1.0000, NS |
|  | Lick | 1.0000, NS |
|  | End-of-trial | 1.0000, NS |

### Nose movement and forces on the head-post during whisking

Head movement, and independently nasal movement, and sniffing have previously been linked to whisking (Towal & Hartmann, 2006; Deschênes, Moore & Kleinfeld, 2012; Moore et al., 2013; Cao et al., 2012; Kurnikova et al., 2017; Dominiak et al., 2019). Here we examined the relationship between whisker motion and nose movement, and forces mice apply on the head post when whisking to touch in our goal directed task. Linear regression models were fit to estimate median nose positions, or median head strain, in each behavioral phase based on the median positions of the four whiskers (Fig 4A). Interestingly, whisker positions were most closely and significantly linked to the nose position immediately after the cue onset -- in the cued-phase (compared to the pre-cue phase; cued, p=0.0000***; trigger, p=0.0745, NS; lick, p=0.6653, NS; end-of-trial, p=1.0000; Wilcoxon signed rank test with Bonferroni correction, N=64 sessions from 14 animals) (Fig 4B).

**Figure 4.**
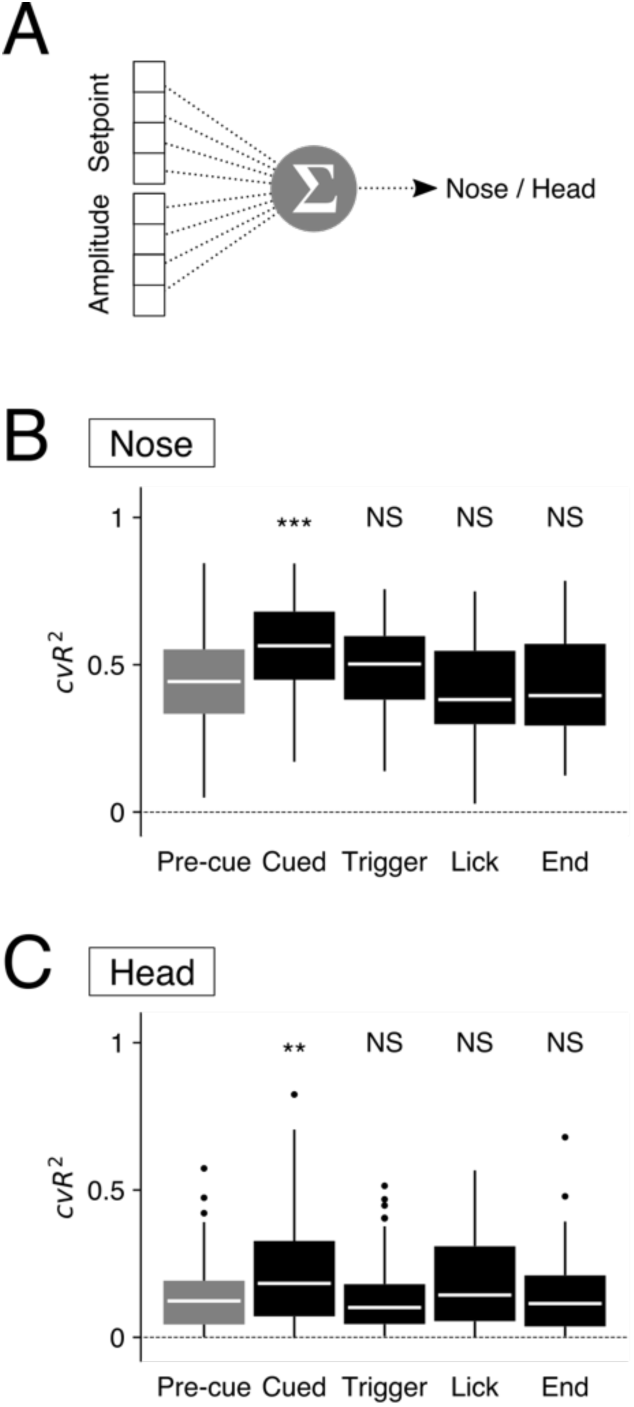
Behavioral phase dependent nose–whisker and head–whisker couplings. A. Model description. The median values during each behavioral time window were computed for the setpoint and amplitude of all the four tracked whiskers. Linear models were fitted to predict the nose position or the head strain during the same behavioral time window based on the eight whisker position related features. Four-fold cross-validation was used to estimate the fitness of the models (cvR^2^). B. Whiskers–nose coupling. The cvR^2^ values are shown in Tukey’s box plots. Without distinction of behavioral time windows (“All”), the linear models explained around the half of the trial-to-trial variability in nose positions. The cvR^2^ values were significantly larger than when the trial-to-trial nose positions were shuffled (“Shuffle”). The linear models restricted to the cued period had significantly larger predictive powers compared to the non-restricted “All” models. On the other hand, the linear models for licking period had significantly smaller predictive powers. *p<0.05, **p<0.01, ***p<0.001, NS, p>0.05, Wilcoxon signed rank test with Bonferroni correction. **N=69 sessions of 15 animals**. C. Whiskers–head coupling. The cvR^2^ values are shown in Tukey’s box plots. Overall predictive power tended to be small but significantly larger than those based on the shuffled data, explaining around 20–30% of the total trial-to-trial variability in head strain values. The linear models based on the triggering time window had significantly smaller predictive power than the non-restricted “All” models. *p<0.05, ***p<0.001, NS, p>0.05, Wilcoxon signed rank test with Bonferroni correction. **N=64 sessions of 14 animals.**

Similarly, whisker position and head post strain were also best linked together after the cue onset (compared to the pre-cue phase; cued, p=0.0086**; trigger, p=1.0000, NS; lick, p=0.5093, NS; end-of-trial, p=1.0000, NS; Wilcoxon signed rank test with Bonferroni correction, N=69 sessions from 15 animals) (Fig 4C). These results imply that the mice recruit nose and head movement to their motor plan when they attempt to move their whiskers to actively touch the contact sensor. The results of this analysis are consistent with the previous studies linking left-right differences in whisking with nose movement (Dominiak et al., 2019).

### Trial-to-trial stereotypy and animal to animal and day to day variability

While we found a few common patterns of whisker/nose movement across behavioral sessions of different animals, we also found a lot of variability across animals, sessions and trials. To examine the source of the variability, and whether this trial-to-trial variability could be attributed to the differences in the strategies that individual mice used to achieve the goal of the task, we used a principal component approach to examine the data. Each trial was represented as a set of features: the median whisking setpoint, median amplitude values, and median nose positions during 3 behavioral phases (the contact trigger, lick and end-of-trial phase). Each trial was visualized using t-SNE which revealed that from session to session and animal to animal, these features could form small clusters (Fig5A, right). The clusters corresponded to trials belonging to a single behavioral session from an animal or to multiple sessions from a single animal (Fig 5A).

**Figure 5.**
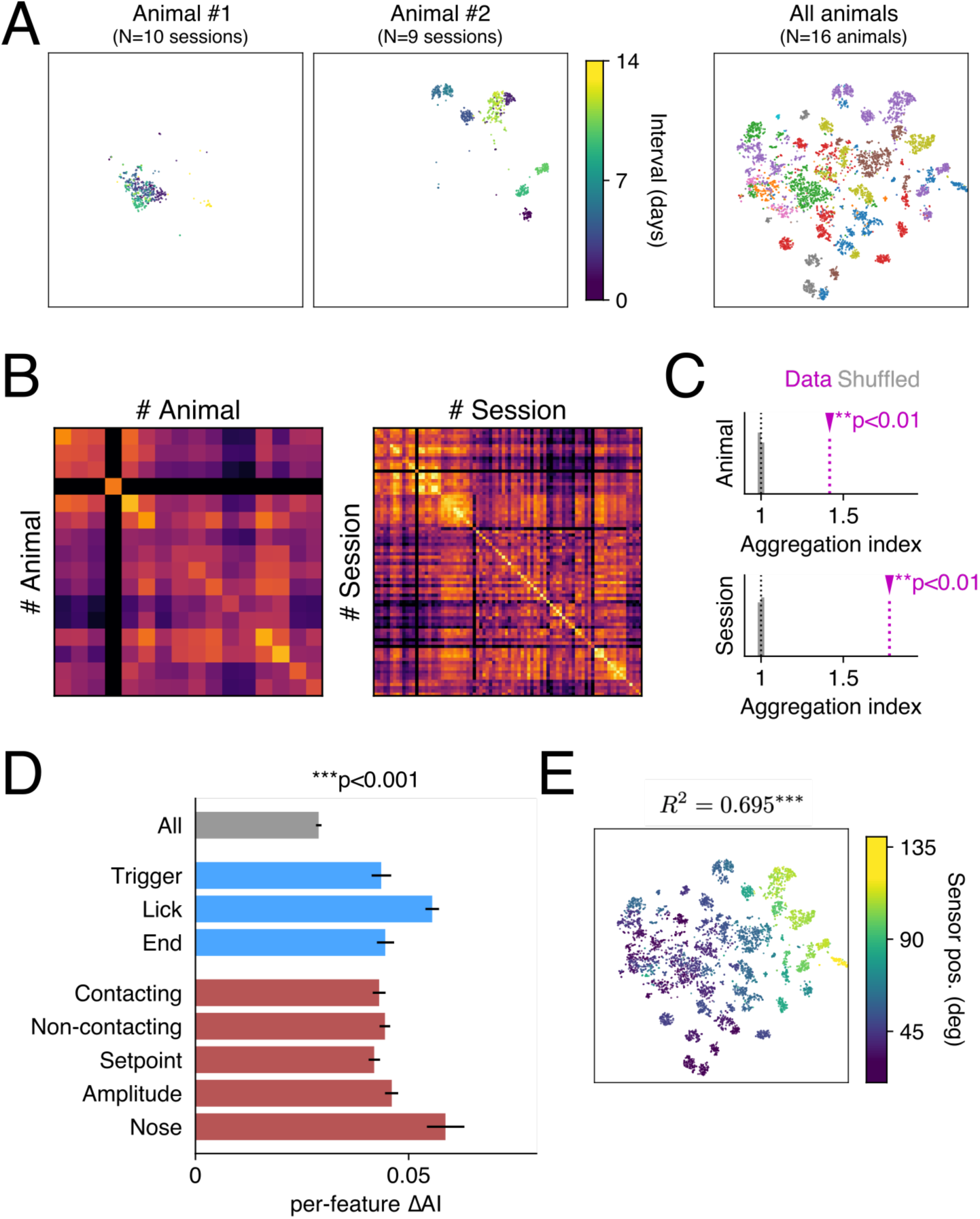
Session-wise stereotypy of the goal-directed behavior of the mice. (A) t-SNE plots of trial-to-trial variability. The left two panels show representative sessions, and the colors representing the interval in days from the earliest session. The rightmost panel shows all the trials from all sessions of all animals recorded. Here, the colors represent individual animals. (B) Analysis of feature-based Euclidian distances. The Euclidean distances were computed based on the Z-scores of individual trials in the high-dimensional feature space. The mean trial-to-trial distance between given two animals (left) or sessions (right) is shown as distance matrices. (C) Aggregation of trial features around individual sessions and animals. Aggregation index (AI) was defined as the ratio of the mean within-cluster trial-to-trial distance to the mean inter-cluster distance. The actual values (magenta) of animal-based (top) and session-based (bottom) AI were significantly larger than the distribution of values based on the shuffled datasets (gray). **p<0.01, based on the comparison of AI values with N=500 shuffled datasets. (D) Contribution of different aspects of the trial to session-wise aggregation. A part of features of trials, related to each of the labels shown, were shuffled, and the corresponding per-feature decreases in session-based AI values from the original dataset were plotted. The bars represent the 95% confidence intervals. Note that the features related to the licking phase, and those related to the nose position, have significantly larger contribution to the observed value of AI. (E) Distribution of session-to-session stereotypy is related to the sensor position. The t-SNE plot same as in (A) is color-coded by the sensor position of the session, in terms of whisking angles of the contacting-side front whisker. The session-median t-SNE coordinates have significant predictive power of the sensor position (***p<0.001, based on the comparison of R^2 values with N=3000 shuffled datasets). **N=16 animals, 85 sessions.**

To examine the state of clustering in detail, we introduced a metric of distance between trials. The extent of the cluster in a behavioral session, within the trial space, was defined as the mean distance between any pair of trials belonging to the session. The distance between a pair of behavioral sessions was defined as the mean distance between any pair of trials out of these two sessions. Animal- or session-based distance matrices revealed a set of bright pixels along the diagonal, indicating measurable similarities between trials belonging to an individual animal or to sessions from a single animal. This tendency appeared clearer for the session-based distance matrix than for the animal-based one (Fig 5B). The aggregation index, i.e. the ratio of inter-cluster distances to the size of clusters (see Materials and Methods for details), was larger for the actual dataset compared to the shuffled datasets, suggesting that clustering arose from the details of movement in trials belonging to single animals or to sessions from individual animals (Fig 5C) (p<0.01, based on comparison with 500 shuffled datasets).

Our definition of trial-to-trial distance included 3 behavioral features: whisking setpoints, whisking amplitudes and nose positions, in the distinct behavioral phases. To determine the contribution of individual behavioral features to trial-to-trial variability, the subset of behavioral features were shuffled, one after another, and the change in the aggregation index was computed (Fig 5D). Our analysis revealed that the animal’s behavior during the licking phase contributed most to the aggregation index compared to the other behavioral phases (p<0.01**, Kruskal–Wallis test and Dunn’s *post hoc* pairwise tests with Bonferroni correction). Furthermore, nose motion had larger per-feature contribution to the aggregation index than the whisker-related features (p<0.01**, Kruskal–Wallis test and Dunn’s *post hoc* pairwise tests with Bonferroni correction).

Together, these results imply that behavioral variability is reflected by movement of individual body parts, during each behavioral phase, and that the movement is linked to “uninstructed” behavior -- nose motion -- not to the goal of the task *per se*, i.e. whisking to touch. Trial-to-trial variability was related to epochs where mice were not engaged in the actual task, i.e. when they were licking.

### Factors mediating behavioral stereotypy and variability

What factors contribute to behavioral variability across animals and sessions? One possibility was that animals use different motor strategies depending on difficulty of the task i.e. on the spatial configuration of the contact sensor and whiskers. To examine this possibility, we tested whether the inter-session distance co-varied with the distance of the contact sensor from the whiskers (Video1, Video2). From session to session, mice could position their whiskers closer (Video1) or farther from the contact sensor (Video2). We found that the *t*-SNE coordinates of individual sessions correlated with the distance of the contact sensor from the resting position of the contacting whisker (R^2^ = 0.6949, linear regression) (Fig 5E). Comparison of the dataset to the 3000 shuffled datasets, where the correspondence between the sensor position and the *t*-SNE coordinates was randomized, revealed that the effect was significant (p<0.001***; shuffled R^2^, min=0.0000, median=0.0169, max=0.1834). This result supports the idea that during cue triggered whisking mice take into account the variability in position of their bodies relative to elements in the environment that they touch. The spatial environment around the face could be one of the most important sources of session-to-session behavioral variability in this task.

## Discussion

Our work shows that mice can control how they move and position individual whiskers. They can control the extent of movement and can even independently control the vigor of whisking on two sides of the face. They can effectively control and coordinate in a behavioral state dependent manner the spread between whiskers. These results suggest an impressive level of fine spatial-temporal control over single facial muscles. In the course of a trial, mice do not just move a single whisker -- a whisker that they have learned to use in a tactile behavior -- but they invariably move whiskers bilaterally and they move their nose side to side and apply up and down forces on the head post. Finally, in a single session mice exhibit an extraordinary level of coordination and stereotypy in moving their whiskers and nose to achieve the goal of touching a sensor. *A priori*, it was possible that from trial to trial and day to day, that mice adjusted how they coordinated their movements, but our work shows the remarkable consistency in what mice move, how they move and how they adapt their movements to small changes in the external environment.

From an anatomic perspective these results are not surprising. The two sides of the face have independent muscles with independent points of attachment on the skull, and each mystacial whisker is associated with a sling muscle (Dörfl, 1982; Wineski, 1983; Haidarliu et al., 2010). It should therefore be possible for rodents to exert control over movement of each mystacial vibrissae independently. Despite the anatomical organization basic questions have lingered: when do rodents control whiskers on just one side of the face? How and when do rodents control single whiskers, and how is this control reflected? Do side to side differences in whisking just reflect direction of head or body movement (Towal & Hartmann, 2006; Dominiak et al., 2019; Bergmann et al., 2022)?

Our work here highlights how anatomy is transformed into function. First, even in highly trained mice, when mice are trained to move a whisker on one side of the face into contact with a sensor, mice invariably move whiskers bilaterally on every trial. This result suggests that facial movements in rodents often have a bilateral component. Second, even though whiskers move bilaterally, movement on the two sides of the face is not identical: whiskers spread apart more on the contacting side, whiskers are positioned distinctly on the two sides, and move with specific vigor at high frequencies on the two sides of the face. Third, mice control the movement of the whisker they used to touch distinctly from the movement of the adjacent whisker. Note that earlier work in head fixed rats using a very similar paradigm, came to the conclusion that rats could move their whiskers independently and that this was mostly evident in the spread between whiskers (Sachdev, Sato & Ebner, 2002). In another study of freely moving rats (Grant et al., 2009), the authors also concluded that rats control movement of individual whiskers and that this was evident in the spread of whiskers. But in their study, whisker spread decreased before contact, before touch. In our hands, mice do something very different -- they increase the spread of whiskers on contacting side. The different results of the two studies are likely to be related to the context. The goal of whisking in our study was for mice to hit a sensor with enough force to get a reward. In the earlier work, rats were not water deprived, there was no task, and rats were simply exploring their environment (Grant et al., 2009). Additionally, in our earlier work, in the course of performing complex task of navigating a plus maze, side to side differences in set point of whisking and movement of the nose were evident on every trial as mice planned and executed turns. Whisking direction predicted movement direction and was coordinated with direction of eye movement (Bergmann et al., 2022; Dominiak et al., 2019). In a different set of studies, in head fixed mice we have shown that the spread between whiskers could be potentially used as a real-time parameter to trigger rewards (Sehara et al., 2021). Taken together these studies highlight varieties of neural-behavioral strategies that rodents use to control movement of individual whiskers bilaterally.

The present work emphasizes that a number of facial movements are coordinated together in the course of mouse actively using a single whisker in a tactile behavior. On one hand it seems that there is complexity and multiple variables to account for -- individual whiskers, nose, head post, i.e. neck muscles. On the other hand, the stereotypy is remarkable: multiple aspects of behavior are coordinated from trial to trial, in some animals even from day to day. A significant portion of the variance in coordination can be explained by small changes in how far mice have to move a whisker.

The coordination and stereotypy in this behavior are remarkable but point to several fundamental issues for systems neuroscientists. First point being emphasized in recent papers by many labs is that variance in neural activity can arise from unexpected sources, like uninstructed movements (Steinmetz et al., 2019; Stringer et al., 2019; Musall et al., 2019). A second point that our work makes is that behavior is fundamentally multimodal involving multiple dimensions that are coordinated together, and that activity of a particular neuron or in a particular circuit can be related to any component of the overall behavior. Dissecting this, understanding what a spike or spikes in a circuit relate to, what triggers the activity or what the activity causes, require additional analytical approaches.

In course of a simple action, the brain or the animal actively controls and coordinates the movement of a number of parts of the body. In the present work, the whisker used to touch is the only instructed movement, all other behaviors -- the movement of adjacent whiskers, the movement of whiskers on the other side of the face, the movement of the nose, and the forces mice apply on the head post -- are uninstructed. Even though it is likely that we have under-sampled the actual number of uninstructed behaviors that occur when mice move a whisker to touch (for example, the posture and limbs probably also change position in a coordinated fashion), it is clear that they move more than just the whisker.

What are the neural mechanisms underlying behavioral variability? One explanation is that variability arises from variability in circuit activity. While it is likely that variability in cortical / subcortical activity contributes to behavior, it is highly unlikely, that individual cortical or subcortical neurons, modulate or coordinate the spectrum of facial movements. What is more likely is that variability in neural activity arises from the interaction between multiple coordinated action sequences -- the behavior -- and the neural circuits that control and respond to the movement. The timing and kinematics of a particular action -- a whisk -- varies because of variability in the other aspects of behavior that modulate activity in the brain.

Understanding how the brain controls active behavior, requires an understanding of the actual details of behavior, the moment-by-moment actions taken to achieve a goal. Our expectation is that knowing what moves when mice move a single whisker to touch will help to better understand both variance in cortical activity and variance in the sensory motor actions mice take in performing tactile behaviors.

## Materials and Methods

### Animals and surgery

We performed all procedures in accordance with protocols for the care and use of laboratory animal approved by the Charité–Universitätsmedizin Berlin and the Berlin Landesamt für Gesundheit und Soziales (LaGeSo). Adult C57bl6 mice (n=16), weighing 25 to 32 g were anesthetized with Ketamine/Xylazine (90mg/kg / 10mg/kg). Lightweight aluminium headposts were affixed to the skull using Rely X and Jet Acrylic (Ortho-Jet) black cement (Ebner et al., 2019; Dominiak et al., 2019). In the two days after surgery, analgesia was provided by Buprenorphine and Carprofen injections.

### Behavioral task

We trained head-fixed mice to use their whiskers to touch a piezo film-based contact sensor. An auditory cue (piezo buzzer) initiated trial and the cue stayed for 2 seconds or until whisker contact with sensor. The contact sensor was placed in front of the C2 whisker of the mouse, with the animal’s whiskers rostral to the C2 being trimmed back to 2 mm. The signal from the contact sensor was fed through a custom-built amplifier, the analogue waveform was thresholded to emit a transistor-transistor logic (TTL) signal. If whisker contact evoked waveform exceeded a threshold, auditory cue was turned off, and reward was delivered. The duration of each trial varied and depended on reaction and movement time after cue onset.

### Behavioral training

One week after surgery, mice were habituated to being handled, and habituated to the behavioral apparatus. In the first days of habituation, mice were acclimated to having their head post handled by the experimenter, and to short (up to several minutes) head fixation. In the course of the first week of habituation, the duration of head fixation was gradually increased from 5 to 40 minutes. In the course of habituation to head fixation, mice were also habituated to having their whiskers painted. During whisker painting, mice were rewarded with sweetened condensed milk.

After a week of habituation, mice were water deprived and were trained to respond to an auditory cue and lick the lick tube. Over the course of 2-3 weeks, mice were introduced to the piezo sensor positioned in front of the painted whiskers on the right side of the face. Initially, the sensor was positioned close to the C2 whisker on the right side, with the whiskers rostral to the C2 trimmed back to 2 mm. During the next training days, the sensor was positioned further way from the mouse. To achieve a consistent success rate for each mouse the position of the sensor varied by a few millimeters from day to day.

### Behavioral tracking

To track whisker and nose positions, behavioral data was recorded at 200 Hz with a Basler acA1920-155uc USB 3.0-camera and a f=25mm / F1.4 objective, being set above the animal. The C1 and C2 whiskers on both sides, as well as the tip of the nose, were painted (UV glow, https://www.uvglow.co.uk/), and illuminated by UV torches. Videos of 200 Hz frame rate were acquired in the proprietary format from Matrox Imaging (https://www.matrox.com/) and later converted into the H.264 format. The acquisition and conversion were accomplished using ZR-view, a custom software (Robert Zollner, Eichenau, Germany). Camera triggers were generated either with the reward trigger, or at the offset of the auditory cue. A five second ring buffer was used to record frames of ±2.5 seconds around the camera triggers.

Whisker and nose positions were estimated from the videos using the custom Python script (videobatch) (https://doi.org/10.5281/zenodo.3407666). The top-view videos first underwent a maximum intensity projection using the videobatch script. Regions of interest (ROI) for tracking were selected manually and separately for the whiskers on both sides of the face and the nose positions, using Fiji’s freehand selection tool (Dominiak et al., 2019; Sehara et al., 2019). Using the Python script, pixels that belonged to a particular hue value were collected and the luma-weighted average position was computed. For frames where the algorithm failed for any reason, values were dropped and were filled in later by linear interpolation.

Licking was monitored with a piezo sensor attached to the lick tube. The signal from the sensor was amplified using a custom-built amplifier and licking waveform was exported to a CED-1401 interface. Head strain was monitored with a strain gauge built into the head post holding-rod. Its signal was amplified using a custom amplifier before being fed into the data acquisition interface.

Behavioral sessions were monitored and recorded using Spike2 (CED, Cambridge, UK) equipped with a Power1401 data acquisition interface. During each session, the duration of the auditory cues, reward triggers, camera triggers and lick event data were acquired, along with analog waveforms of the output of the contact sensor, piezo-based lick sensor and strain gauge. The acquired data was saved as compressed Spike2 files and exported as tab-separated value (TSV) files for analysis.

### General analytical procedures

Analytical procedures were performed using Python (https://www.python.org/, version 3.7.6), along with several standard modules for scientific data analysis (NumPy, https://www.numpy.org/, version 1.18.1; Scipy, https://www.scipy.org/, version 1.4.1; matplotlib, https://matplotlib.org/, 3.1.3; pandas, https://pandas.pydata.org/, 1.0.1; scikit-learn, https://scikit-learn.org/stable/, version 0.22.1; Bottleneck, https://pypi.org/project/Bottleneck/, 1.3.2; Statsmodels, https://www.statsmodels.org/, version 0.13.0; h5py, https://www.h5py.org/, version 3.6.0) and non-standard packages (sliding1d, https://github.com/gwappa/python-sliding1d, 1.0; fitting2d, https://doi.org/10.5281/zenodo.3782790, 1.0.0a2). Kruskal–Wallis test and Dunn’s *post hoc* test was performed using a custom python script based on the one found at https://gist.github.com/alimuldal/fbb19b73fa25423f02e8. Data was sorted based on trial duration. Trials on which mice failed to contact the sensor in 2 seconds were not analyzed.

### Behavioral data preprocessing

The whisker positions tracked during each behavioral session were converted to whisking angles. Using the fitting2d Python library, a circle was fit to the set of two-dimensional positions for each whisker, and the position at each timepoint was converted into the polar coordinates around the fitted circle. Whisking setpoint and amplitude were then estimated based on these whisking angles. Upper and lower envelopes of whisker positions were estimated by smoothing the whisker positions using sliding-window mean of a 300 ms time window. The lower envelope was defined as the setpoint; the difference between the upper and lower bounds of the envelope was defined as the whisking amplitude. The mode of whisking setpoints was estimated from the histogram of whisking offsets during each behavioral session. The lower half (up to 50 percentile) of the histogram was fitted using a cubic spline, and the mode was defined as the whisker position giving the maximal density of setpoint positions.

Unless otherwise indicated, this mode of whisking setpoints was considered as the resting position of the whisker and was used as the zero-position of the whisking angles. In the particular cases when the resting positions of individual whiskers were compared, whisking angles were calculated relative to the position of the eyes. The corners of the left and right eyes were annotated manually on a video frame of each behavioral session. A line connecting the corners of the two eyes was drawn. The crossing point between this line and the circle of whisking was defined as the eye level which was then used to calculate the whisking angles.

The position of the contact sensor relative to the mouse could change slightly from day to day, as each animal was positioned anew in the behavioral rig. The sensor position for each day / each session was measured post hoc using the maximal density of whisker positions evident in the video data. Nose position data were projected on a 1D axis of the first principal component.

### Analysis of behavioral epochs

Data from each trial was first parsed into behavioral epochs: a pre-cue phase where mice wait for cue onset, an auditory cue phase which initiated whisker movement, a trigger epoch where whisker contact with the sensor crossed threshold including 500 ms before touch and 250 ms after touch, a lick/reward epoch, and a post-reward epoch (Table 1).

To examine nose-whisker and head-whisker coordination, linear regression models were used to predict the nose position and the head strain based on whisker dynamics. In doing so, time points belonging to a single behavioral phase of all the trials during a single behavioral session were pooled. The values of whisker setpoints and whisking amplitude were used to predict the nose position and the head strain on the same time point. The models were fit using the least-square approach. Four-fold cross-validation scheme was used – the linear model was trained based on 75% of the data, and prediction accuracy was tested on the remaining 25% based on the coefficient of determination (cross-validated R^2^, or cvR^2^) -- and the mean of the four cvR^2^ scores was used as the cvR^2^ value of the given session.

To estimate trial-to-trial variability, we used 9 features of behavior tracked in 3 behavioral epochs (trigger / lick / end-of-phase) to generate a 27-dimensional vector. The 9 features consisted of: (a) the median whisking offsets of the four tracked whiskers, (b) the median whisking amplitudes of the four tracked whiskers, and (c) the median nose position. Trial-to-trial variability was mapped using two types of methods. The *t*-SNE method was used to visualize distribution of individual trials in a 2-dimensional space (van der Maaten and Hinton, 2008), whereas the Euclidean method was used to compute trial-to-trial distances. Before applying the Euclidean method, all of the 27 features were standardized by computing the Z scores, i.e. subtracting their means and then dividing it by their standard deviations. The Euclidean distance in the 27-dimensional space of Z scores was used as the distance metric between trials.

Euclidean trial-to-trial distances were used to estimate the diameters of groups (single animals and single sessions), and inter-group distances. The diameter of the group of trials belonging to a single animal or a session was determined as the mean distance between two randomly chosen trials from the group. The inter-group distance was determined as the mean distance between two randomly chosen pair of trials, each of which from one from the pair of groups. Because it was computationally demanding to compute distances across all the >4000 trials, we randomly sampled 100,000 pairs of trials and computed their inter-trial distances.

To examine the state of clustering in accordance with groups of trials, we devised the aggregation index (AI), as the ratio of the mean inter-group distance (i.e. mean trial-to-trial distance between different groups) to the mean group diameter (i.e. mean trial-to-trial distance within a single group).

To relate trial-to-trial variability to the position of the contact sensor, we computed 2-D coordinates of each behavioral session in the *t*-SNE space as the median of all the trials belonging to the session. A linear regression model was trained using the least-squares approach, to predict the sensor position during each session based on the 2-D coordinates of the session.

### Sample size and statistics

For the analysis of whisker and nose movements, we developed scripts for automated tracking of orofacial -- whiskers, nose -- movement (Dominiak et al., 2019; Bergmann et al., 2021). Two criteria were used to select sessions: there had to be more than 10 trials in a session and success rate in each session had to be more than 50%. As a result, for analysis related to Figures 3-5, we had data for 82-85 sessions from 16 animals in total. In one behavioral session nose positions were not tracked and this data was excluded from the analysis. For the analysis of whisking setpoints and frequency power, manual annotation of eye position was needed, and we completed annotation for 48 behavioral sessions in 7 animals. Comparisons were made using Wilcoxon’s signed-rank test.

For the analysis of whisker and nose movements, two criteria were used to select sessions: there had to be more than 10 trials in a session and success rate in each session had to be more than 50%. As a result, we used 85 sessions from 16 animals in total. In one behavioral session nose positions were not tracked and this data was excluded from the analysis.

For each trial, behavioral phases were used only when the whole duration of the phase was recorded in the 5 second video. Unless otherwise noted, comparisons were made using Kruskal–Wallis test, followed by Dunn’s *post hoc* pairwise tests with Bonferroni correction. The analysis of trial-to-trial variability was based on three epochs: trigger onset, licking and end-of-trial phases. The data related to these epochs could be easily extracted from video.

To statistically compare whether the given aggregation index was significantly higher than the chance level, we prepared 500 different datasets where annotation of each trial to the group (animal or session) was shuffled. All the resulting aggregation indices were represented as the deviation from the median value of the shuffled data, and the p-value was computed based on where the index of the true data resides relative to the other shuffled data.

To examine the contribution of different groups of parameters to trial-to-trial variability, we prepared 500 datasets (per group) where the given group of parameters were shuffled. The resulting reduction in the aggregation index (ΔAI) was considered to be the contribution of the shuffled parameters to trial-to-trial variability. Comparisons between shuffled data were performed using Kruskal–Wallis test, followed by Dunn’s *post-hoc* pairwise tests with Bonferroni correction.

To examine statistical significance in the correlation between trial-to-trial variability and the contact sensor position, the R^2^ value of the linear regression model was used. Apart from generating a model based on the actual dataset (N=84 sessions), 3000 models based on random datasets were prepared and the correspondence between the shuffled 2-D session coordinates and the sensor position were examined. Statistical testing was based on comparison of the R^2^ value of the model of the actual data and those of the 3000 shuffled models; the R^2^ values were sorted based on the distance from the median value of the shuffled models, and the p-value for the R^2^ of the true dataset was computed by assessing what percentile the value was located.

## Acknowledgements

We would like to thank the Charite workshop in particular Jan-Erik Ode, Alexander Schill and Daniel Deblitz for machine and electronics work.

## Author contributions

RS and KS planned and designed the analysis and experiments and wrote the manuscript. MS, NB, and SD performed the experiments. MS, KS, NB analyzed the data. MEL designed experiments and wrote the manuscript.

## References

Adibi, M. (2019) Whisker-Mediated Touch System in Rodents: From Neuron to Behavior. Frontiers in systems neuroscience. 13. doi:10.3389/FNSYS.2019.00040.

Arkley, K., Grant, R.A., Mitchinson, B. & Prescott, T.J. (2014) Strategy change in vibrissal active sensing during rat locomotion. Current biology:24 (13), 1507–1512. doi:10.1016/J.CUB.2014.05.036.

Avitan, L. & Stringer, C. (2022) Not so spontaneous: Multi-dimensional representations of behaviors and context in sensory areas. Neuron: 110: 3064–3075 10.1016/j.neuron.2022.06.019.

Berg, R.W. & Kleinfeld, D. (2003) Rhythmic whisking by rat: retraction as well as protraction of the vibrissae is under active muscular control. Journal of neurophysiology. 89: 104–117. doi:10.1152/JN.00600.2002.

Bergmann, R., Sehara, K., Dominiak, S.E., Kremkow, J., Larkum, M.E. & Sachdev, R.N.S. (2022) Coordination between eye movement and whisking in head-fixed mice Navigating a Plus Maze. eNeuro. 9, ENEURO.0089-22.2022. doi:10.1523/ENEURO.0089-22.2022.

Cao, Y., Roy, S., Sachdev, R.N.S. & Heck, D.H. (2012) Dynamic correlation between whisking and breathing rhythms in mice. Journal of Neuroscience. 32: 1653–1659. doi:10.1523/JNEUROSCI.4395-11.2012.

Carvell, G.E. & Simons, D.J. (1990) Biometric analyses of vibrissal tactile discrimination in the rat. Journal of Neuroscience. 10: 2638–2648. doi:10.1523/JNEUROSCI.10-08-02638.1990.

Churchland, M.M., Afshar, A. & Shenoy, K. V. (2006) A central source of movement variability. Neuron. 52: 1085–1096. doi:10.1016/J.NEURON.2006.10.034.

Deschênes, M., Moore, J. & Kleinfeld, D. (2012) Sniffing and whisking in rodents. Current Opinion in Neurobiology. 22: 243–250. doi:10.1016/J.CONB.2011.11.013.

Dolensek, N., Gehrlach, D.A., Klein, A.S. & Gogolla, N. (2020) Facial expressions of emotion states and their neuronal correlates in mice. Science. 368: 89–94. (6486). doi:10.1126/SCIENCE.AAZ9468.

Dominiak, S.E., Nashaat, M.A., Sehara, K., Oraby, H., Larkum, M.E. & Sachdev, R.N.S. (2019) Whisking asymmetry signals motor preparation and the behavioral state of Mice. Journal of Neuroscience. 39: 9818–9830. doi:10.1523/JNEUROSCI.1809-19.2019.

Dörfl, J. (1982) The musculature of the mystacial vibrissae of the white mouse. Journal of Anatomy. 135: 147–154.

Ebner, C., Ledderose, J., Zolnik, T.A., Dominiak, S.E., Turko, P., Papoutsi, A., Poirazi, P., Eickholt, B.J., Vida, I., Larkum, M.E. & Sachdev, R.N.S. (2019) Optically induced calcium-dependent gene activation and labeling of active neurons using CaMPARI and Cal-Light. Frontiers in Synaptic Neuroscience. doi:10.3389/FNSYN.2019.00016.

Finlayson, K., Lampe, J.F., Hintze, S., Würbel, H. & Melotti, L. (2016) Facial indicators of positive emotions in Rats. PloS One.11 doi:10.1371/JOURNAL.PONE.0166446.

Ghazanfar, A.A. & Schroeder, C.E. (2006) Is neocortex essentially multisensory? Trends in Cognitive Sciences. 10: 278–285. doi:10.1016/J.TICS.2006.04.008.

Grant, R.A., Mitchinson, B., Fox, C.W. & Prescott, T.J. (2009) Active touch sensing in the rat: anticipatory and regulatory control of whisker movements during surface exploration. Journal of neurophysiology. 101 (2), 862–874. doi:10.1152/JN.90783.2008.

Grant, R.A., Mitchinson, B. & Prescott, T.J. (2012) The development of whisker control in rats in relation to locomotion. Developmental psychobiology. 54 (2), 151–168. doi:10.1002/DEV.20591.

Grant, R.A., Sperber, A.L. & Prescott, T.J. (2012) The role of orienting in vibrissal touch sensing. Frontiers in Behavioral Neuroscience. 6. doi:10.3389/FNBEH.2012.00039.

Haidarliu, S., Golomb, D., Kleinfeld, D. & Ahissar, E. (2012) Dorsorostral snout muscles in the rat subserve coordinated movement for whisking and sniffing. Anatomical Record. 295:1181–1191. doi:10.1002/AR.22501.

Haidarliu, S., Kleinfeld, D., Deschênes, M. & Ahissar, E. (2015) The musculature that drives active touch by vibrissae and nose in mice. Anatomical Record. 298: 1347–1358. doi:10.1002/AR.23102.

Haidarliu, S., Simony, E., Golomb, D. & Ahissar, E. (2010) Muscle architecture in the mystacial pad of the rat. The Anatomical Record: Advances in Integrative Anatomy and Evolutionary Biology. 293: 1192–1206. doi:10.1002/ar.21156.

Hartmann, M.J. (2001) Active sensing capabilities of the Rat whisker system. Autonomous Robots 2001 11:3. 11 (3), 249–254. doi:10.1023/A:1012439023425.

Hartmann, M.J.Z. (2011) A night in the life of a rat: vibrissal mechanics and tactile exploration. Annals of the New York Academy of Sciences. 1225: 110–118. doi:10.1111/J.1749-6632.2011.06007.X.

Kelso, J.A.S. (2009) Synergies: Atoms of brain and behavior. In: D. Sternad (ed.). *Progress in Motor Control. Advances in Experimental Medicine and Biology*. Boston, MA, Springer. pp. 83–91. doi:10.1007/978-0-387-77064-2_5.

Knutsen, P.M., Derdikman, D. & Ahissar, E. (2005) Tracking whisker and head movements in unrestrained behaving rodents. Journal of Neurophysiology. 93: 2294–2301. doi:10.1152/JN.00718.2004.

Knutsen, P.M., Pietr, M. & Ahissar, E. (2006) Haptic object localization in the vibrissal system: behavior and performance. Journal of Neuroscience. 26: 8451–8454. doi:10.1523/JNEUROSCI.1516-06.2006.

Krakauer, J.W., Ghazanfar, A.A., Gomez-Marin, A., MacIver, M.A. & Poeppel, D. (2017) Neuroscience needs behavior: Correcting a reductionist bias. Neuron. 93:480–490. doi:10.1016/J.NEURON.2016.12.041.

Kurnikova, A., Moore, J.D., Liao, S.M., Deschênes, M. & Kleinfeld, D. (2017) Coordination of orofacial motor actions into exploratory behavior by rat. Current Biology. 27: 688– 696. doi:10.1016/J.CUB.2017.01.013.

Langford, D.J., Bailey, A.L., Chanda, M.L., Clarke, S.E., Drummond, T.E., et al. (2010) Coding of facial expressions of pain in the laboratory mouse. Nature methods. 7: 447–449. doi:10.1038/NMETH.1455.

McElvain, L.E., Friedman, B., Karten, H.J., Svoboda, K., Wang, F., Deschênes, M. & Kleinfeld, D. (2018) Circuits in the rodent brainstem that control whisking in concert with other orofacial motor actions. Neuroscience. 368: 152–170. doi:10.1016/J.NEUROSCIENCE.2017.08.034.

Moore, J.D., Deschênes, M., Furuta, T., Huber, D., Smear, M.C., Demers, M. & Kleinfeld, D. (2013) Hierarchy of orofacial rhythms revealed through whisking and breathing. Nature. 497: 205–210. doi:10.1038/NATURE12076.

Moore, J.D., Kleinfeld, D. & Wang, F. (2014) How the brainstem controls orofacial behaviors comprised of rhythmic actions. Trends in Neurosciences. 37: 370–380. doi:10.1016/J.TINS.2014.05.001.

Murakami, M. & Mainen, Z.F. (2015) Preparing and selecting actions with neural populations: toward cortical circuit mechanisms. Current Opinion in Neurobiology. 33: 40–46. doi:10.1016/J.CONB.2015.01.005.

Musall, S., Kaufman, M.T., Juavinett, A.L., Gluf, S. & Churchland, A.K. (2019) Single-trial neural dynamics are dominated by richly varied movements. Nature Neuroscience. 22: 1677–1686. doi:10.1038/S41593-019-0502-4.

Sachdev, R.N.S., Sato, T. & Ebner, F.F. (2002) Divergent movement of adjacent whiskers. Journal of Neurophysiology. 87: 1440–1448. doi:10.1152/JN.00539.2001.

Saraf-Sinik, I., Assa, E. & Ahissar, E. (2015) Motion makes sense: an adaptive motor-sensory strategy underlies the perception of object location in rats. Journal of Neuroscience. 35: 8777–8789. doi:10.1523/JNEUROSCI.4149-14.2015.

Schroeder, J.B. & Ritt, J.T. (2016) Selection of head and whisker coordination strategies during goal-oriented active touch. Journal of Neurophysiology. 115: 1797–1809. doi:10.1152/JN.00465.2015.

Sehara, K., Bahr, V., Mitchinson, B., Pearson, M.J., Larkum, M.E. & Sachdev, R.N.S. (2019) Fast, flexible closed-loop feedback: Tracking movement in “Real-millisecond-time”. eNeuro. 6 (6). doi:10.1523/ENEURO.0147-19.2019.

Sehara, K., Zimmer-Harwood, P., Larkum, M.E. & Sachdev, R.N.S. (2021) Real-time closed-loop feedback in behavioral time scales using deeplabcut. eNeuro. 8 (2). doi:10.1523/ENEURO.0415-20.2021.

Severson, K.S., Xu, D., van de Loo, M., Bai, L., Ginty, D.D. & O’Connor, D.H. (2017) Active touch and self-motion encoding by merkel cell-associated afferents. Neuron. 94: 666–676.e9. doi:10.1016/J.NEURON.2017.03.045.

Severson, K.S., Xu, D., Yang, H. & O’Connor, D.H. (2019) Coding of whisker motion across the mouse face. eLife. 8. doi:10.7554/ELIFE.41535.

Steinmetz, N.A., Zatka-Haas, P., Carandini, M. & Harris, K.D. (2019) Distributed coding of choice, action and engagement across the mouse brain. Nature. 576: 266–273. doi:10.1038/S41586-019-1787-X.

Stringer, C., Pachitariu, M., Steinmetz, N., Reddy, C.B., Carandini, M. & Harris, K.D. (2019) Spontaneous behaviors drive multidimensional, brainwide activity. Science. 364:255. doi:10.1126/SCIENCE.AAV7893.

Tantirigama, M.L.S., Zolnik, T., Judkewitz, B., Larkum, M.E. & Sachdev, R.N.S. (2020) Perspective on the multiple pathways to changing brain states. Frontiers in Systems Neuroscience. 14. doi:10.3389/FNSYS.2020.00023.

Towal, R.B. & Hartmann, M.J. (2006) Right-left asymmetries in the whisking behavior of rats anticipate head movements. Journal of Neuroscience. 26: 8838–8846. doi:10.1523/JNEUROSCI.0581-06.2006.

van der Loos, H. & Woolsey, T.A. (1973) Somatosensory cortex: structural alterations following early injury to sense organs. Science. 179: 395–398. doi:10.1126/SCIENCE.179.4071.395.

van der Maaten L., Hinton G. (2008) Visualizing data using t-SNE. J Machine Learning Res 9(86):2579−2605.

Vincent, S.B. (1912) The functions of the vibrissae in the behavior of the white rat. Chicago, IL, University of Chicago.

Waschke, L., Kloosterman, N.A., Obleser, J. & Garrett, D.D. (2021) Behavior needs neural variability. Neuron. 109: 751–766. doi:10.1016/J.NEURON.2021.01.023.

Welker, W.I. (1964) Analysis of sniffing of the Albino rat. Behaviour. 22: 223–244. doi:10.1163/156853964X00030.

Wineski, L.E. (1985) Facial morphology and vibrissal movement in the golden hamster. Journal of Morphology. 183: 199–217. doi:10.1002/jmor.1051830208.

Wineski, L.E. (1983) Movements of the cranial vibrissae in the Golden hamster (*Mesocricetus auratus*). Journal of Zoology. 200: 261–280. doi:10.1111/j.1469-7998.1983.tb05788.x.

Zagha, E., Erlich, J.C., Lee, S., Lur, G., O’Connor, D.H., Steinmetz, N.A., Stringer, C. & Yang, H. (2022) The importance of accounting for movement when relating neuronal activity to sensory and cognitive processes. The Journal of Neuroscience. 42: 1375– 1382. doi:10.1523/JNEUROSCI.1919-21.2021.

